# HLA Type and Chronic Viral Infection Impact Peripheral T-cell Receptor Sharing Between Unrelated Individuals

**DOI:** 10.1101/2021.03.19.436220

**Authors:** Sarah A. Johnson, Spencer L. Seale, Rachel M. Gittelman, Julie A. Rytlewski, Harlan S. Robins, Paul A. Fields

## Abstract

The human adaptive immune system must generate extraordinary diversity to be able to respond to all possible pathogens. The T-cell repertoire derives this high diversity through somatic recombination of the T-cell receptor (TCR) locus, a random process that results in repertoires that are largely private to each individual. However, certain factors such as low junctional diversity, thymic selection, and T-cell proliferation upon antigen exposure can affect TCR sharing among individuals. By immunosequencing the TCRβ variable region of 426 healthy individuals, we find that fewer than 1% of TCRβ clones are shared between individuals on average, consistent with largely private TCRβ repertoires. However, we detect a significant correlation between increased HLA allele sharing and increased number of shared TCRβ clones, with each additional shared HLA allele contributing to an increase in ∼0.01% of the total TCRβ clones being shared, supporting a key role for HLA type in shaping the immune repertoire. Surprisingly, we find that shared antigen exposure to CMV leads to fewer shared TCRβ clones, even after controlling for HLA, indicative of a largely private response to major viral antigenic exposure. Consistent with this hypothesis, we find that increased age is correlated with decreased overall TCRβ clone sharing, indicating that the pattern of private TCRβ clonal expansion is a general feature of the T-cell response to other infectious antigens. All of these factors contribute to shaping the TCRβ repertoire, and understanding their interplay has important implications for the use of T cells for therapeutics and diagnostics.

## INTRODUCTION

T cells make up a key component of the adaptive immune response and allow the body to respond to the diverse range of pathogens it may encounter. The adaptive immune system of a healthy adult includes up to 10^15^ highly diverse T cells (1,2). Antigen recognition depends on both T-cell specificity and the molecular complex presenting the antigen. Foreign antigens are first processed and presented by an individual’s major histocompatibility complex (MHC). T cells that encounter their specific cognate MHC-presented antigen will bind and proliferate, leading to an immune response. The vast diversity of possible T-cell receptors (TCR) is generated by the random recombination of genes in the third complimentary determining regions (CDR3) within a TCR’s α and β chains. In the recombination process of the β chain, loci of the variable (V), diversity (D), and joining (J) regions are randomly spliced together with non-templated insertions and deletions occurring between each junction, resulting in up to 10^11^ possible sequences (3,4). Recombination of the TCRα chain includes only V and J gene segments, resulting in fewer possible rearrangements and making the TCRβ chain a more suitable target for identifying unique T cells, and thus the focus of this paper. T cells mature in the thymus, where their affinity to MHC molecules is tested prior to subsequent release into the periphery. Successful antigen recognition requires T cells to effectively recognize the body’s MHC and coordinate a response. However, excessive avidity to the MHC causes T cells to incorrectly identify host cells as foreign targets and may result in autoimmunity. Therefore, in healthy individuals self-immunogenic T cells are targeted for apoptosis, while those yielding mild affinity to the MHC are released into the periphery for circulation (5). As the MHC is encoded by highly polymorphic human leukocyte antigen (HLA) loci in humans, this process of thymic selection occurs within the context of an individual’s HLA type. As a result, VDJ recombination, HLA restriction, and antigen exposure collectively contribute to a largely private TCRβ repertoire.

Despite the large space of potential TCRβ rearrangements, the existence of public clones found in two or more individuals has been well characterized, and occurs more frequently than would be expected by chance (6–8). Public TCRβ clones can arise through either convergent recombination - due to highly probable rearrangements, or convergent selection - due to proliferation after common antigen exposure. Biases in VDJ recombination and junctional indel patterns have been computationally modeled, and suggest that CDR3 sequences that more closely resemble the germline-encoded nucleotide sequence are more likely to occur and thus be shared between individuals (9,10). Previous studies have identified publicly expanded, antigen-specific T cells to a variety of pathogens including CMV and SARS-CoV-2, among others (11–14), supporting convergent selection as a mechanism for generating public clones. Together, both of these processes contribute to the existence of public clones, but much remains to be discovered about how factors such as additional antigen exposure and HLA type modify them.

Understanding the forces underlying the inherent diversity of the TCRβ repertoire of an individual and the public sharing of TCRβ clones is of great clinical interest. Decreased diversity of TCRβ clones in older individuals has been associated with reduced immune function (15). Similarly, increased evolutionary divergence of HLA class I alleles has been correlated with better responses to immune checkpoint inhibitors in cancer patients (16). Public TCRβs that respond to specific antigens have the potential to be used diagnostically to identify an individual’s antigen exposure (11–13). Likewise, antigen-specific clones have demonstrated therapeutic potential as next-generation CAR-T cells (17,18). Additionally, HLA-restriction of TCRβs is clinically relevant to evaluating histocompatibility for the purposes of bone-marrow and solid organ transplants (19,20). Continued development of such immunological and medical advancements depends on fully understanding the determinants that shape public versus private TCRβs.

To explore the influence of biological and environmental forces on the dynamics of TCRβ clone sharing, we utilized a published set of TCRβ repertoires from 426 healthy human subjects (8,11). We assessed the role of HLA zygosity, HLA allele sharing, CMV exposure, and age in shaping the immune repertoire and the sharing of TCRβ clones between individuals. By analyzing both the highest-frequency and single-copy TCRβ clones, we identified a consistent positive association between numbers of shared HLA alleles and TCRβ clones. Additionally, we found that CMV exposure and increased age result in more private TCRβ repertoires, in particular among high frequency clones. Our results demonstrate the impact of both age and HLA type on TCRβ clone generation and maturation, influencing the sharing of low frequency clones. In contrast, our results indicate that infectious antigen exposure leads to the expansion of largely private and HLA-restricted TCRβ clones, and that it impacts the sharing of high frequency clones.

## RESULTS

### Determinants of TCRβ Repertoire Diversity in Healthy Individuals

We first investigated how HLA type influences diversity of the TCRβ repertoire. The divergent allele hypothesis suggests that greater diversity of HLA alleles leads to a greater diversity of presented peptides (21,22), and thus potentially greater diversity of the TCRβ repertoire. To address this question, we utilized previously published TCRβ repertoire data from over 600 individuals with known HLA type and CMV serostatus. We restricted analysis to individuals with full 4-digit resolution at 6 HLA loci (HLA-A, HLA-B, HLA-C, HLA-DRB1, HLA-DQB1, HLA-DPA1), for which > 90% of individuals in the cohort have a resolved type. To control for any technical variation due differences in T-cell fraction or input material, we additionally restricted analysis to only those 426 subjects with greater than 200,000 total T cells. We determined the zygosity of these individuals at the included HLA loci and quantified repertoire diversity with two metrics: richness and clonality. The richness of each repertoire was calculated as the number of unique TCRβ nucleotide rearrangements after computationally downsampling all repertoires to 200,000 productive templates. Simpson clonality was also calculated for each repertoire to assess the dominance of high frequency clones in the repertoire. Within this cohort, 44% of individuals are homozygous at the HLA-DPA1 loci, and heterozygous at all other loci (S1 Fig). In contrast to our expectations, there was not a significant relationship between the number of homozygous class I or class II HLA alleles and the richness or clonality of an individual’s repertoire in this cohort (Figs 1A, 1B, S2A, S2B). Similarly, there is not an overall correlation across all 6 included loci (S2C, S2D Figs). This suggests that, in addition to not influencing the overall richness, HLA zygosity does not influence the extent of oligo-clonal dominance in the repertoire.

**Figure 1.**
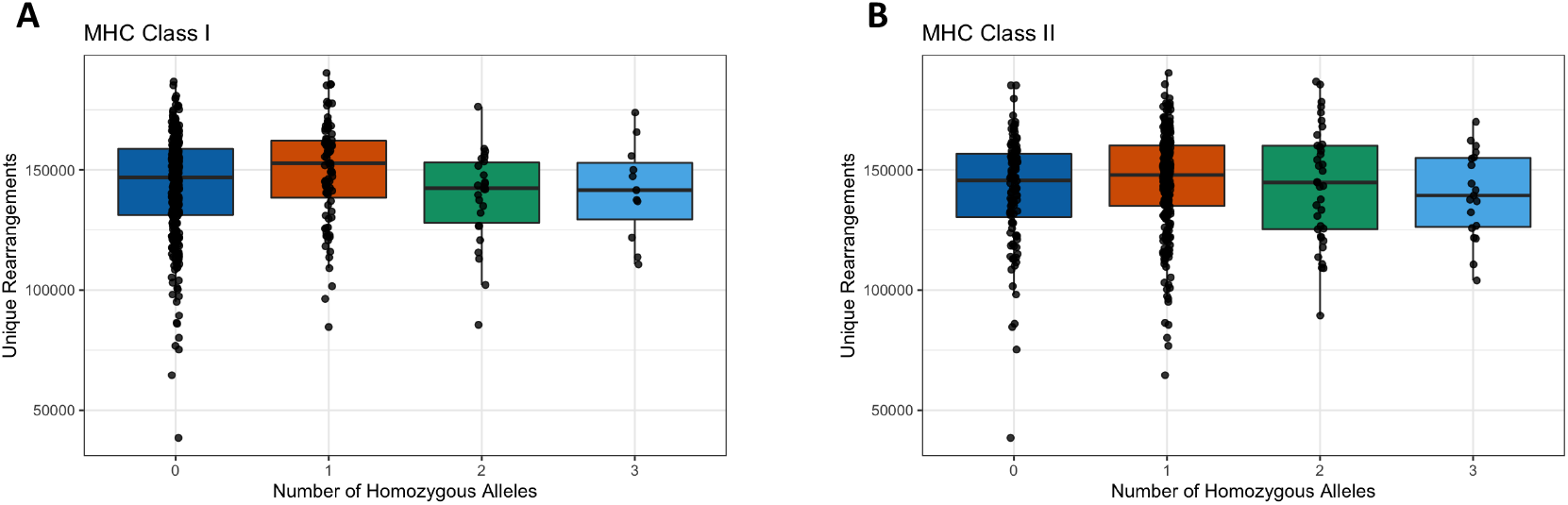
Correlation of TCRβ repertoire richness and HLA zygosity. TCRβ richness was quantified as the number of unique nucleotide rearrangements in a repertoire computationally downsampled to 200,000 productive templates. HLA zygosity at neither (A) class I loci nor (B) class II loci correlated with diversity (Spearman rho = 0.035, p = 0.47 and rho = 0.021, p = 0.66 respectively).

We additionally examined the influence of age and CMV exposure on the diversity of the TCRβ repertoire. Consistent with prior studies, we found that CMV+ individuals have increased clonality and decreased repertoire richness, indicating a more focused repertoire post-viral exposure (S3A, S3B Figs), and also observed a negative correlation between age and repertoire diversity (S3C, S3D Figs) (23–25). Within our cohort, we did not see a correlation between overall HLA zygosity and diversity among either CMV+ or CMV-individuals (S2E and S2F Figs). Taken together, this data demonstrates that the diversity of a person’s TCRβ repertoire is largely independent of HLA zygosity and may be driven more by age and exposure to antigens.

### Healthy Individuals with More Shared HLA Alleles Share More TCRβ Clones

While HLA zygosity does not affect the overall diversity of a TCRβ repertoire, positive thymic selection does select for TCRβ clones with affinity to an individual’s specific set of HLA alleles (26). We thus hypothesized that shared HLA alleles between unrelated individuals may contribute to the sharing of unique clones. The number of shared clones between each pairwise combination of repertoires was determined by comparing TCRβ clones by their functional identity (the V gene family, CDR3 amino acid sequence, and J gene). This allows clone sharing to be detected when different individuals generate TCRβ clones with the same specificity but distinct nucleotide sequences through VDJ recombination. To be considered a shared TCRβ clone, an exact match was required. The HLA allele sharing between individuals was determined regardless of HLA zygosity, where two individuals homozygous for the same HLA allele were considered to share two alleles.

We saw a significant correlation between the number of HLA alleles shared between two individuals and the number of shared unique downsampled TCRβ clones (p < 0.001, Mantel test, Fig 2A). Using a linear mixed-effects model, individuals with no HLA alleles in common shared on average 1,884 of their 200,000 downsampled clones (0.94%), while each additional shared HLA allele resulted in an increase of 14 shared clones. This suggests a baseline population of public clones found across all subjects regardless of HLA type, consistent with the existance of bystander public clones with high generation probability, in addition to a subset of clones that are shared based on HLA type (10,27). While there was a similar relationship between shared alleles and shared TCRβ clones within both class I and class II alleles, this correlation was slightly stronger for HLAII alleles (Figs 2B and 2C). This is consistent with CD4+ T cells, which bind class II alleles, making up a greater fraction of the repertoire than CD8 + T cells, which bind class I alleles (28).

**Figure 2.**
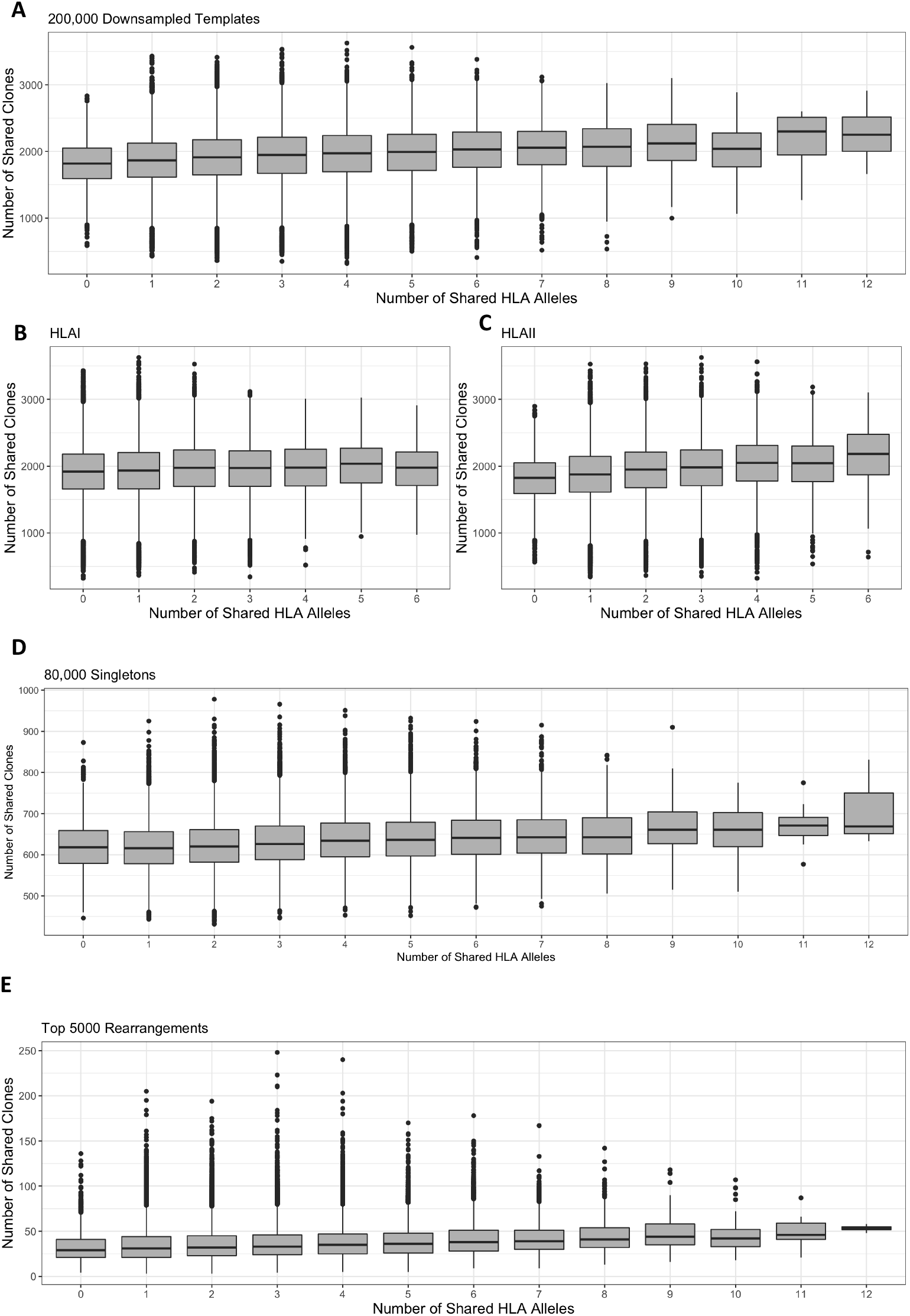
TCRβ clone sharing is correlated with HLA allele sharing. Among repertoires computationally downsampled to 200,000 productive templates, clone sharing positively correlated with increased numbers of (A) shared HLA alleles overall, and of both HLA (B) class I and (C) class II alleles (A: Mantel rho = 0.17, p < 1e-3, linear mixed effects model slope = 14, intercept = 1884. B: Mantel rho = 0.07, p < 1e-3, linear mixed effects model slope = 9, intercept = 1920. C: Mantel rho = 0.19, p < 1e-3, linear mixed effects model slope = 21, intercept = 1881). (D) The number of singletons shared between two individuals was positively and significantly correlated with the number of HLA alleles shared (Mantel rho = 0.14, p < 1e-3, linear mixed effects model slope = 6, intercept = 613). (E) The number of clones shared among the top 5,000 unique rearrangements was positively and significantly correlated with the number of HLA alleles shared between two individuals (Mantel rho = 0.13, p < 1e-3, linear mixed effects model slope = 1.3, intercept = 34).

### Sharing of the Low Frequency and Expanded TCRβ Clones is Impacted by HLA Type

We next wanted to characterize the extent to which the number of shared HLA alleles relates to the sharing of both expanded and low frequency clones. While immunosequencing cannot distinguish naïve vs. memory T cells, we hypothesized that many singletons (TCRβ clones seen once in a sample) correspond to naïve clones, while the highest frequency rearrangements correspond to clones that have likely expanded in response to prior antigen exposure. To characterize the singleton repertoire, we randomly selected 80,000 singletons from the repertoire of each individual. Singleton clones were selected by nucleotide sequence to best capture rearrangements that were present once in the full repertoire of an individual, but compared between individuals using functional identity. While this process may exclude highly-probable rearrangements that are present multiple times in an individual (29), it captures the lowest frequency clones in a repertoire. Similar to what we observed when analyzing the downsampled repertoires, there was a significant correlation between the number of shared HLA alleles and number of shared singletons between individuals (p < 0.001, Mantel test, Fig 2D). Notably, individuals with no shared alleles still shared on average 613 of 80,000 (0.77%) singletons. Since T cells that are present once in a repertoire are less likely to have encountered antigens, this overlap of singletons between individuals may be attributed to convergent recombination of frequently generated TCRβ rearrangements. Each shared HLA allele was correlated with an increase of 6 shared singletons, consistent with a role for HLA type in the maturation of the TCRβ repertoire. However, even when individuals shared 10 HLA alleles, their repertoires were still largely private, sharing on average 660 of 80,000 singletons, still < 1%. Thus, while HLA type plays an important role in the maturation of T cells, the random VDJ somatic recombination results in most individuals having a highly private TCRβ repertoire.

To characterize the expanded repertoire of each subject, we selected the top 5,000 unique clones by frequency from each full repertoire prior to downsampling. As chronic infections such as CMV increase the clonality of TCRβ repertoires (11,30) (S3A Fig), we selected a common number of rearrangements to mitigate the effect of a few largely expanded clones. Similar to the results among the downsampled and singleton repertoires, there was a significant relationship between the number of HLA alleles shared and number of high-frequency clones shared (p < 0.001, Mantel test, Fig 2E). On average, individuals that shared no HLA alleles shared ∼34 of their top 5,000 clones (0.68%), with each additional shared allele leading to an additional 1.3 shared clones. Together, these findings suggest that while the HLA type of an individual does not impact the overall diversity of their TCRβ repertoire, HLA type does have a small but significant impact on the specific TCRβ clones selected for maturation, as well as those that expand in response to antigens.

### Shared HLA Alleles Increases Sharing of Rare TCRβ Clones

Given the role of convergent recombination in influencing TCRβ clone sharing, we next sought to assess whether HLA type influences sharing of clones with high-generation probability differently than that of those with lower generation probability. To do this, we determined the generation probabilities of all TCRβ rearrangements using OLGA (9), which correlated strongly with CDR3 length and the number of indels (S4A-C Figs). We established a cutoff point (P_gen_ = 1.66e-09) to distinguish rare from common TCRβ clones by finding the intersection point of generation probabilities between public TCRβ clones that occurred in more than one subject and private TCRβ clones found in only one subject (Fig 3A, S4D Fig). Each TCRβ clone was thus classified as “rare,” with generation probabilities lower than this cutoff, or “common” otherwise. Sharing of each subset of clones between individuals was examined in both the downsampled and singleton repertoires. While sharing of both common and rare clones were significantly correlated with increased sharing of HLA alleles, the relationship was stronger among rare clones (Mantel rho 0.28) compared to common clones (Mantel rho 0.12) (p < 0.001, Mantel test, Figs 3B and 3C). As rare TCRβ clones are those with rearrangements not commonly generated during VDJ recombination, these results suggest that their selection for maturation may be modulated by HLA genotype.

**Figure 3.**
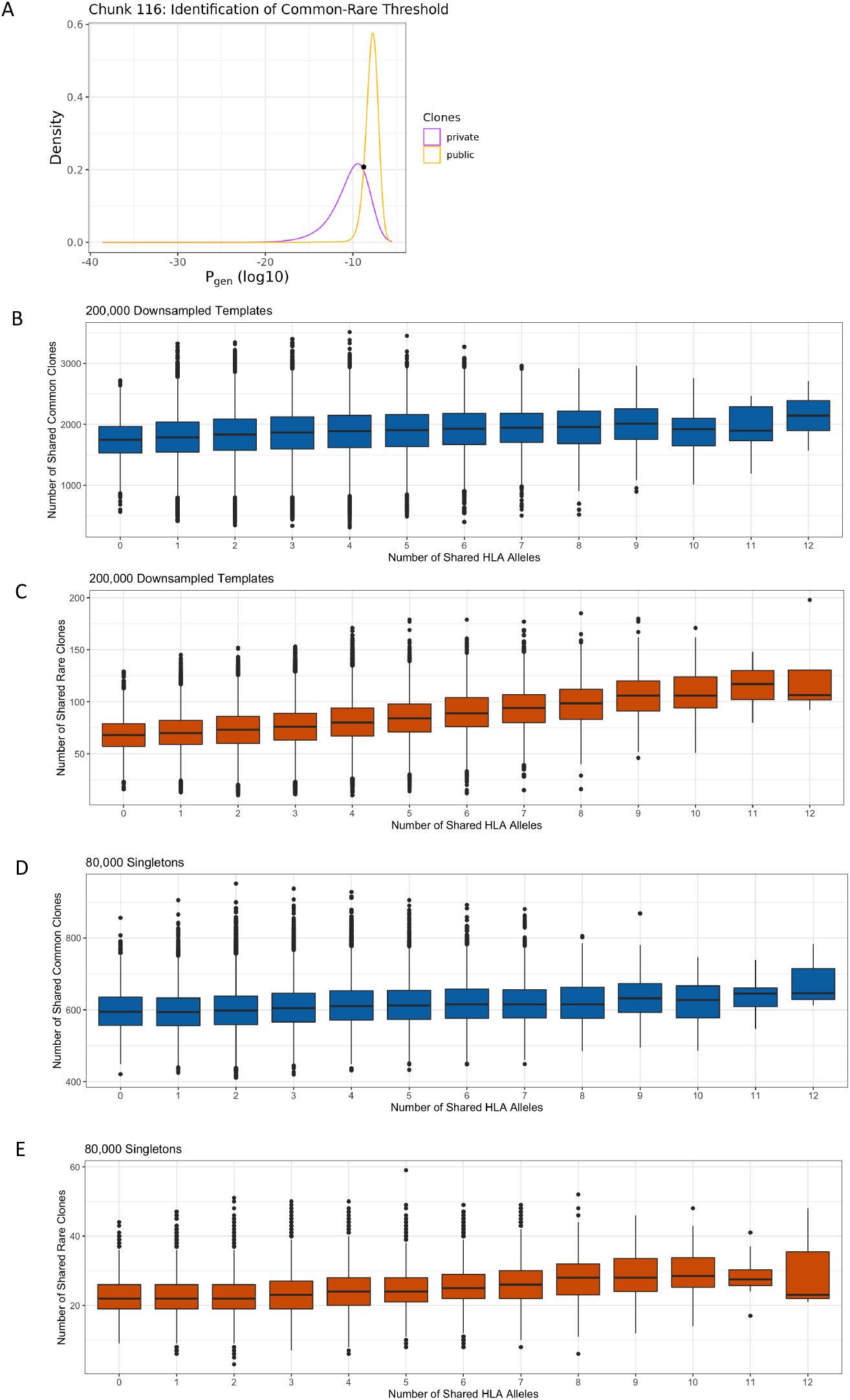
TCRβ clone sharing among common and rare clones. Clones were classified as common versus rare by identifying a generation probability cutoff point between clones occurring once and those occurring more than once in 31 different “chunks” representing the entirety of TCRβ rearrangements observed in the downsampled data. (A) Intersection point from a single representative “chunk.” The number of shared HLA alleles was significantly correlated with clone sharing among both (B) common (Mantel rho = 0.12, p < 1e-3) and (C) rare clones (Mantel rho = 0.28, p < 1e-3). Similarly, the number of shared HLA alleles was significantly correlated with sharing of singletons among (D) common (Mantel rho = 0.14, p < 1e-3) and (E) rare (Mantel rho = 0.18, p < 1e-3) clones.

Interestingly, and in contrast to the downsampled repertoires, the singleton TCRβ repertoires exhibited similar positive associations between HLA allele sharing and both shared common (Mantel rho = 0.14) and rare (Mantel rho = 0.18) clones (p < 0.001, Mantel test, Figs 3D and 3E). These results suggest that HLA allele sharing affects both rare and common TCRβ clone sharing, but the degree of impact depends on the frequencies of those clones within an individual repertoire. TCRβ repertoire frequency is largely determined by antigen exposure, suggesting that expansion of rare clones may increase clone sharing among individuals with matched HLA types, though these clones only make up a small proportion of the overall repertoire.

### CMV Negative Individuals Share More TCRβ Clones Than CMV Positive Individuals

Our prior analyses focused on clone sharing independent of antigen exposure. We next sought to determine whether common viral antigenic exposure would lead to differences in clone sharing. We focused our analysis on CMV, as our analysis cohort was split roughly equally between CMV+ (subjects with past CMV infection) and CMV-individuals (11). We hypothesized that CMV+ individuals may share more clones than CMV-individuals. Surprisingly, we observed the opposite, with CMV-individuals sharing more downsampled TCRβ clones than CMV+ subjects (Fig 4A), and significantly more clones than a permuted null distribution (S5 Fig; permuted P = 0.002). The pattern of increased clone sharing between CMV-individuals persisted among both rare and common TCRβ clones, regardless of the number of HLA clones shared (S6A and S6B Figs). This suggests that CMV infection leads to significantly fewer shared clones between individuals.

**Figure 4.**
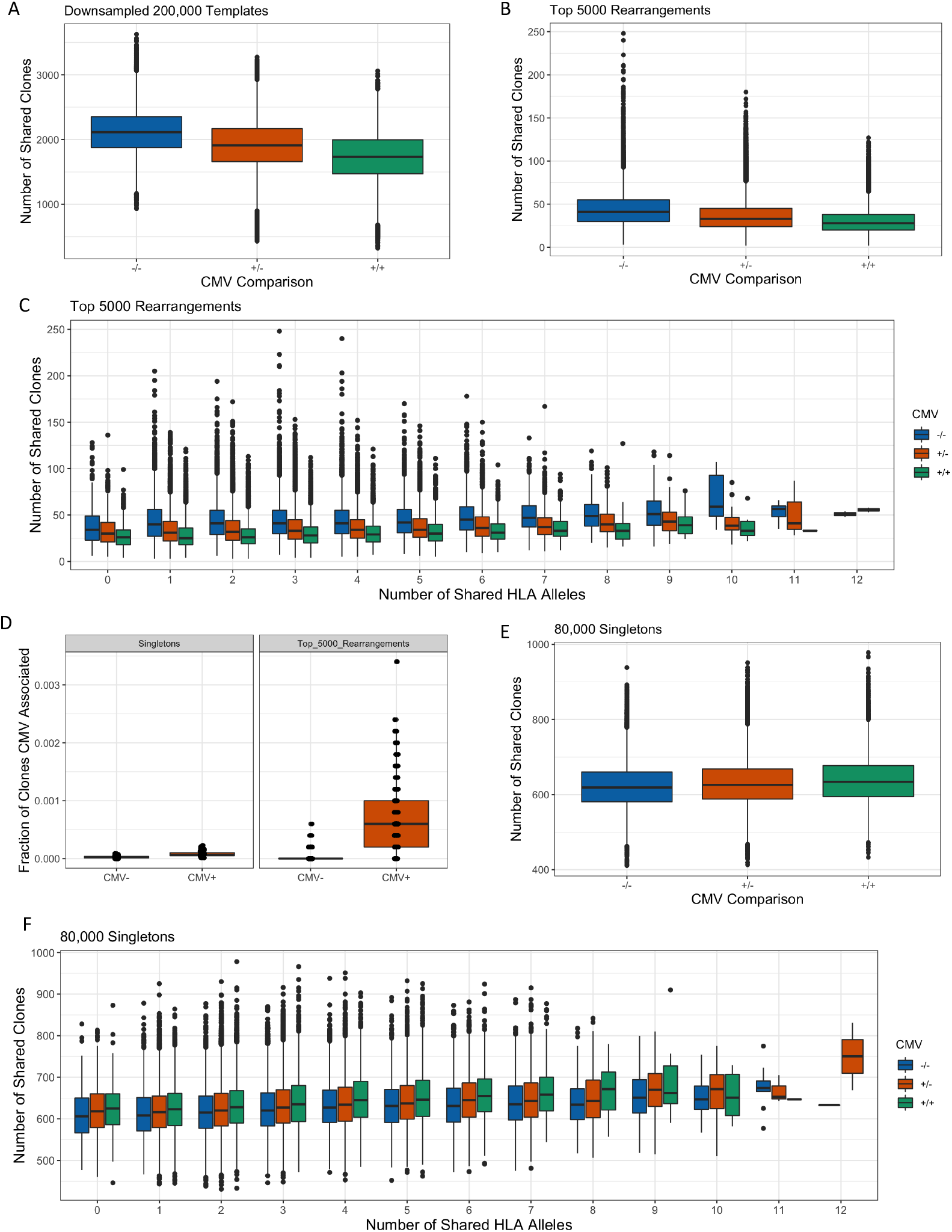
CMV status influences TCRβ clone sharing. (A) CMV-individuals shared more TCRβ clones within their downsampled repertoires compared to CMV+ individuals. (B) CMV-individuals shared more of their top 5,000 rearrangements compared to CMV+ individuals. (C) CMV-individuals shared more top rearrangements than CMV+ individuals do, regardless of the number of HLA alleles shared. (D) There was a greater fraction of clones in CMV+ individuals that were among the 164 CMV associated clones identified in Emerson et al. compared to CMV-individuals, among both singletons and top 5,000 rearrangements (Wilcoxon rank sum tests, p < 2.2e-16 for both). (E) There was a similar number of singletons shared among CMV- and CMV+ individuals. (F) CMV+ individuals with a common number of shared HLA alleles shared slightly more singletons than CMV-individuals do.

CMV+ individuals have significantly higher clonality and lower richness than CMV-individuals (S3A and S3B Figs), effectively decreasing the number of unique clones that could be shared in CMV+ individuals (25). To account for this disparity in diversity, we again looked at clone sharing among the top 5,000 unique TCRβ rearrangements. There was also greater sharing of top TCRβ clones between CMV-individuals compared to CMV+ individuals (Fig 4B) and compared to the null (permuted P = 0.002). This suggests that TCRβ repertoires become more private after CMV antigen exposure, likely through the expansion of mostly private clones. Notably, this trend is consistent independent of shared HLA alleles (Fig 4C). Despite the overall decrease in shared TCRβ clones compared to CMV-individuals, CMV+ individuals showed significantly greater enrichment in their top 5,000 clones of CMV-associated TCRβ clones, as identified in Emerson et al. (Fig 4D, P <2.2e-16, Unpaired Wilcox test). This suggests that while some public clones expand based on common viral antigen exposure, most of the expanded antigen-specific clones are private to a given repertoire.

In contrast to the most expanded clones, individuals shared a comparable number of singletons regardless of CMV status (Fig 4E). We did observe a significant enrichment of CMV-associated clones among the singletons of CMV+ individuals (Fig 4D, P <2.2e-16, Unpaired Wilcox test), which may reflect some antigen exposed TCRβ clones among the singletons. When stratified by shared HLA alleles, CMV+ individuals share slightly more singletons than CMV-individuals (Fig 4F), which can likely be explained by differences in age between CMV+ and CMV-individuals. We observed that CMV+ individuals in this cohort tend to be older than CMV-individuals (S3E Fig), and that older individuals had a greater proportion of common singletons than younger individuals (S7C and S8C Figs), making increased singleton sharing between CMV+ individuals consistent with the less diverse and more public singleton repertoires of older individuals (15).

### TCRβ Repertoires Become Increasingly Private with Age

Given the association between CMV and TCRβ clone sharing, we hypothesized that antigen exposure in general may lead to more private TCRβ repertoires. Older individuals are likely to have encountered more antigens over the course of their life, consistent with more clonal repertoires in older subjects (S3F Fig) (31). To examine whether age, as a proxy for antigen exposure, is a contributing factor in clone sharing, we divided individuals into above/equal to or below the median age (42 years) and compared TCRβ clone sharing within both the downsampled and the singleton repertoires. We found that older individuals shared fewer downsampled clones than younger individuals (median 1842 vs. 2136) (Fig 5A), and significantly fewer clones than the null (permuted P = 0.002). Since CMV+ individuals tend to be older (S3E Fig), we also looked at the impact of age within CMV+ and CMV-individuals. Older individuals continued to share fewer clones independent of CMV status (Fig 5B). Additionally, clone sharing between younger individuals continued to be greater than clone sharing between older individuals regardless of the number of HLA alleles shared (Fig 5C). This suggests that exposure to more pathogenic antigens, such as CMV, results in expansion of a set of largely private TCRβ clones.

**Figure 5.**
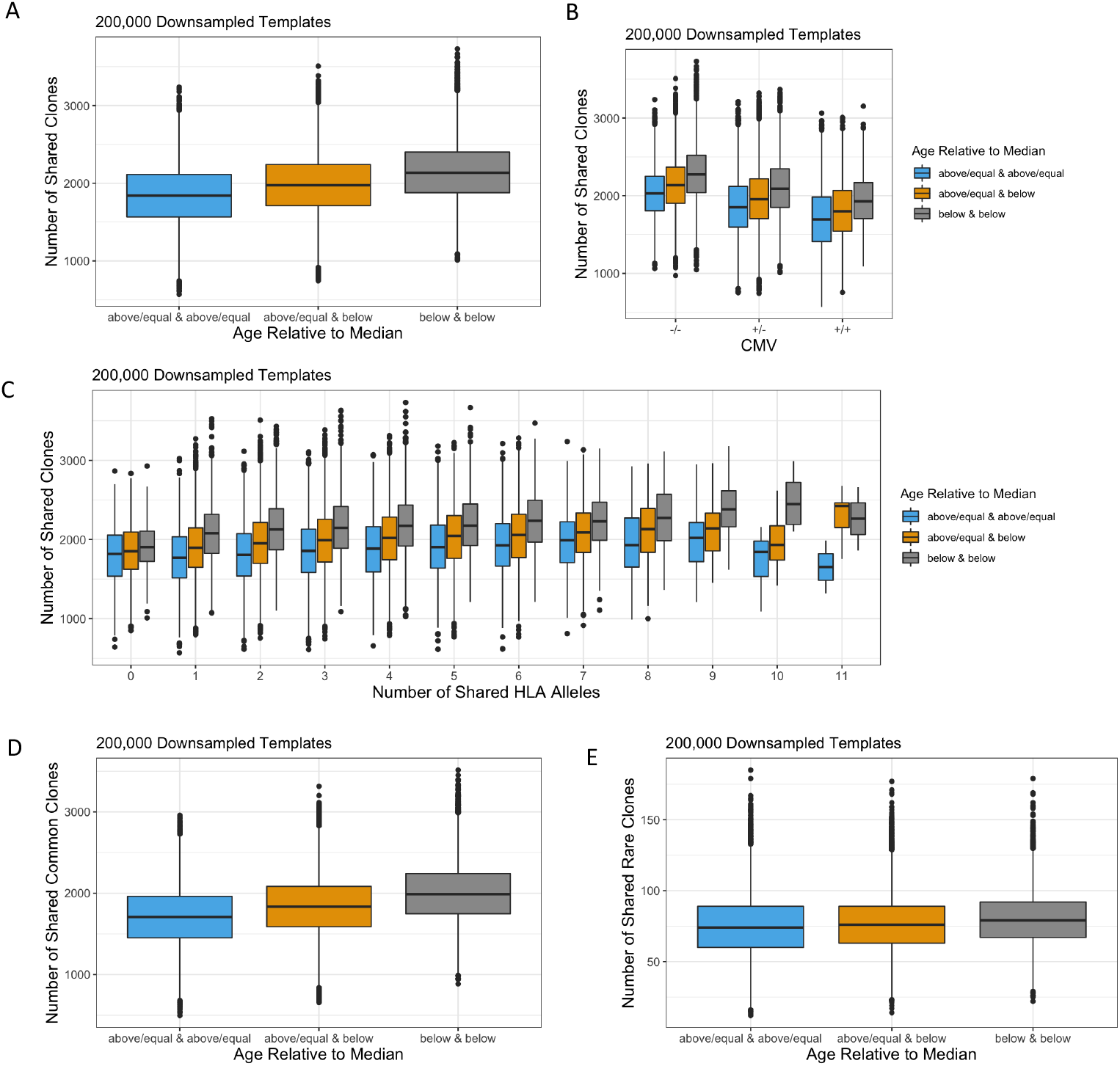
Impact of age on TCRβ clone sharing. Individuals were stratified by the median age (42). (A) Younger individuals shared more clones in their downsampled repertoires compared to older individuals, independent of (B) CMV exposure and (C) shared HLA alleles. Among (D) common and (E) rare clones, younger individuals shared more clones in their downsampled repertoires than older individuals.

To evaluate whether age impacts shared clones differently based on their generative probability, we again split up clonotypes into rare and common subsets using the previously defined cutoff (Fig 3A, S4D Fig). Older individuals shared fewer common clones than younger individuals (median 1707 vs. 1989) (Fig 5D), and significantly fewer common clones than the null (permuted P = 0.002). Older individuals additionally shared fewer rare clones than younger individuals (median 74 vs 79) (Fig 5E), although not significantly fewer rare clones than the null (permuted P > 0 .05). This demonstrates that older individuals share fewer TCRβ clones independent of generation probability.

### Older Individuals Share More Singletons Than Younger Individuals

Next, we examined how age shapes the sharing of singletons. In contrast to the downsampled repertoire, older individuals shared more singletons than younger individuals (median 634 vs 617) (Fig 6A), and significantly more clones than the null (permuted P < 0.05). This trend held true regardless of the number of HLA alleles shared (Fig 6B). Additionally, older individuals shared more singletons in both the rare and the common subsets (Figs 6C and 6D), and significantly more rare and common singletons than the null (permuted (rare) P < 0.05 and (common) P = 0.002). While the expansion of unique clones in response to antigen exposure can explain the decreased sharing in the downsampled repertoire of older individuals, greater TCRβ overlap within the singleton repertoire of older individuals is consistent with the diminished TCRβ diversity in aging immune systems caused by decreased thymopoiesis (32,33). This is furthermore supported by the proportion of rare and common clones within older and younger individuals. Younger and older individuals have a comparable proportion of rare clones among their downsampled repertoire (p = 0.5, Wilcox test) (S8A Fig). However, among the 5000 most abundant TCRβ rearrangements, age was significantly correlated with lower generation probabilities (S7B Fig, Spearman rho = -0.37, p = 8.6e-15) and a higher proportion of rare TCRβ clones (S8B Fig, p = 3.6e-10, Wilcox test). This suggests that antigen-specific TCRβ clones, including those with low generation probability, proliferate due to antigen exposure and thus are enriched in older individuals. In contrast, median generation probabilities of singletons increased significantly with age (S7C Fig, Spearman rho = 0.17, p = 0.0015), and older individuals had a significantly lower proportion of rare singletons compared to younger individuals (S8C Fig, p = 0.03, Wilcox test). These age-dependent differences within the singleton repertoire support previous reports that a reduction in the thymus’s production of naïve TCRβ clones diminishes diversity within this subset, and can explain the greater overlap of singletons among older individuals. Together, these findings support a model whereby antigen exposure to pathogens such as CMV during the life of an individual leads to expansion of mostly rare, private clones, while age related changes in the thymus reduce the complexity and diversity of singletons among older individuals.

**Figure 6.**
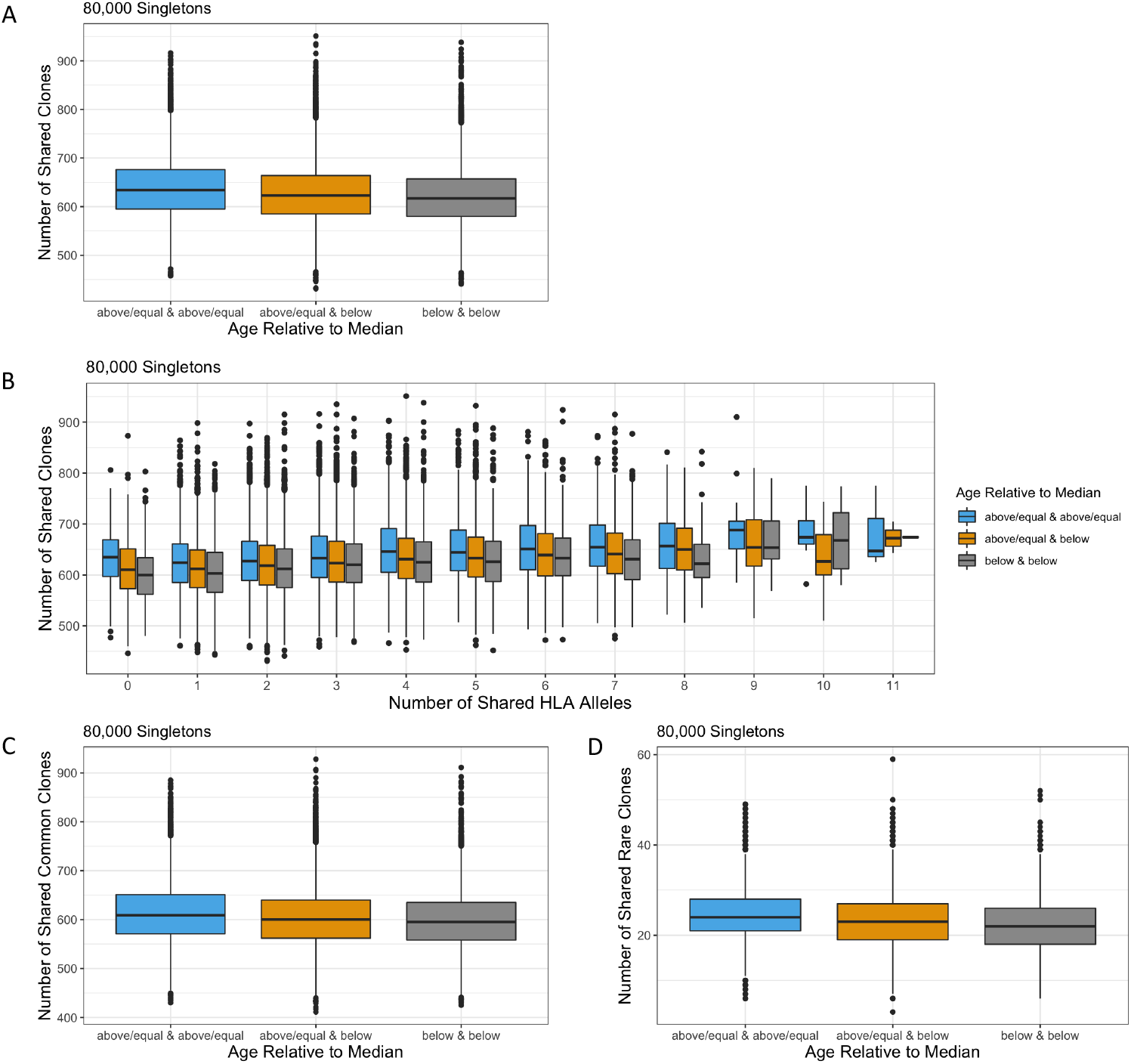
Impact of age on sharing of singleton TCRβ clones. Individuals were stratified by the median age (42). (A) Older individuals shared more singletons than younger individuals. (B) Age impacted the sharing of singletons regardless of the number of HLA alleles. Older individuals shared more (C) common and (D) rare singletons than younger individuals.

## DISCUSSION

The composition of an individual’s TCRβ repertoire is the result of many forces, both biological and environmental (34). While several of these factors have been well characterized, their interaction within the context of HLA type has not been thoroughly explored. Here we looked at the extent to which HLA type, antigen exposure, and age impact the sharing of TCRβ clones between unrelated individuals. We assessed these factors within the context of TCRβ generation probabilities to determine how clones of varying prevalence are affected. By separately surveying clone sharing among singletons and high frequency clones, we distinguished between HLA-restricted selection vs. HLA-restricted expansion. Our data support a broad model of TCRβ proliferation wherein HLA allele sharing increases sharing of TCRβ clones across all frequencies, while increased age correlates with a more public naïve TCRβ repertoire and antigenic exposure results in a more private expanded TCRβ repertoire.

Consistent with previous work characterizing the diversity of the immune repertoire (10,11,14), our data reinforces that the TCRβ repertoire of an individual is predominantly private. Even when several HLA alleles are shared between two unrelated individuals, we see that over 98% of each repertoire is comprised of unique clones. Within the singleton repertoire, which best reflects the inherent diversity of the naïve repertoire, less than 1% of the repertoire is shared between individuals, demonstrating the high variability of VDJ somatic recombination. We additionally show that HLA zygosity of an individual does not strongly affect the overall diversity of their TCRβ repertoire. While previous work with a similar dataset found a correlation between HLA class I zygosity and decreased repertoire richness among CMV-individuals, that analysis quantified diversity with different metrics and did not control for sampling depth, which could impact the finding (30). One limitation of this cohort is that over half of the individuals are of European descent, and a more diverse cohort may have a different distribution of HLA zygosities. Nonetheless, these findings suggest that the space of potential TCRβ clones is diverse enough that the homozygosity of a person’s HLA genotype does not affect the overall diversity of their repertoire.

The divergent allele advantage hypothesis suggests that more heterozygous HLA loci and greater evolutionarily divergence among HLA alleles leads to an increased number of antigens that can be presented to T cells by the MHC (21). It has been further shown that even within fully heterozygous individuals, evolutionary divergence of HLA alleles is positively correlated with the number of peptides predicted to be recognized by the TCRβ repertoire of an individual (16,22). As such, it is perhaps not surprising that HLA zygosity alone is not enough to alter the shape of the TCRβ repertoire. Future work incorporating phylogeny-aware or distance-based clustering of HLA alleles (16,35,36) could better assess the functional diversity of an individual’s HLA alleles and thus the relationship between potential bound peptides and diversity. However, it is also important to note that decreased numbers of bound peptides may not translate to a decrease in the number of unique T cells generated, as HLA restriction during T-cell maturation functions independently of antigen exposure.

Despite the large inherent diversity of the TCRβ repertoire, comparisons between unrelated individuals always contain a portion of public clones. We show that individuals sharing more HLA alleles share more TCRβ clones, suggesting that HLA type impacts the specific T cells selected for maturation and consistent with HLA restriction during thymic selection. Previous work has also shown that public clones occur more frequently than would be expected by chance due to biases in VDJ recombination and indels (1,6,10,37). Our analyses support these findings, as individuals sharing no HLA alleles still share TCRβ clones. Moreover, we see a stronger correlation between HLA sharing and the sharing of rare clones compared to the sharing of common clones, suggesting a background of easily generated clones while rare clones are more HLA restricted. Notably, even among commonly recombined clones, HLA restriction is significantly correlated with clone sharing, suggesting that convergent recombination alone cannot explain the prevalence of all public clones.

While high frequency clones may be shared between individuals after exposure to a common antigen, the lowest frequency clones within an individual should be largely antigen naïve. We demonstrate that the sharing of singleton clones between individuals is a function of both an individual’s HLA genotype and age, consistent with previously observed patterns of HLA restriction and thymic involution, respectively. We see that the number of shared HLA alleles is positively correlated with the number of shared singletons between individuals, further supporting our assertion that sharing of TCRβ clones of all frequencies is influenced by HLA allele overlap. Additionally, our data showing that older individuals share more singleton clones than younger individuals is consistent with thymic involution resulting in decreased T-cell diversity (32). As such, our work highlights the importance of incorporating HLA genotype and age in models examining public clone sharing, as well as the important distinctions between the naïve and memory compartments. This work further supports the hypothesis that TCRβ repertoires can be used to infer the HLA type of an individual (11), however age may be a confounding factor and including younger individuals in the training data may increase model specificity and sensitivity.

Finally, while there has been demonstrated success in the diagnostic potential of utilizing public antigen-specific clones, our work suggests a largely private expansion of TCRβ clones in response to CMV antigen exposure. While public TCRβ clones do exist, they may represent the minority of the overall response. We show that CMV exposure actually decreases overall clone sharing, regardless of the number of HLA alleles shared. This suggests that within a similar HLA context, clonal expansion after CMV exposure is largely private. By extending this analysis to include the age of each individual, as a proxy for continued pathogenic exposure, our data indicate that the expansion of private clones in response to antigen exposure is not unique to CMV. Older individuals have likely been exposed to a broader range of pathogens than younger individuals, shaping their TCRβ repertoires in an increasingly private manner as T cells specific to encountered antigens expand and remain in the memory compartment. Interestingly, work in identical twins has similarly shown predominantly private responses to common antigens across individuals with the same genetic background (38–40). These data underscore how finding public, antigen-specific clones is a difficult problem requiring large datasets. However, the individual nature of TCRβ repertoires provides utility in tracking clones within individuals over time or between individuals with allogeneic T-cell transplants. While posing a considerable computational challenge, important future work will include defining motifs or implementing clustering algorithms to identify TCRβ clones that bind the same antigen (36,41). Together, this work emphasizes the inherent diversity and private nature of human TCRβ repertoires, as well as the importance of incorporating HLA genotypes into models predicting both public TCRβ sharing and antigen-specific expansion.

## METHODS

### Sample Details

All samples analyzed in this study were previously published in Emerson et al. (11). Of the 666 immunosequenced healthy adult PBMC repertoires, we included 426 individuals in this analysis. These individuals all have complete 4-digit HLA resolution at the 6 included HLA loci (HLA-A, HLA-B, HLA-C, HLA-DPA, HLA-DQB, HLA-DRB), CMV serotyping, and 200,000 or more productive templates sequenced. HLA typing and CMV serostatus testing was conducted at the time of sample collection. Additionally, 15 samples were removed due to unexpected sharing of high-frequency TCRβ clones. Of these 426 samples, we know the age at collection of 366 of the subjects who were thus included in the age-related analyses. Samples were computationally downsampled to 200,000 productive nucleotide templates with replacement to minimize skewing in the number of singletons. TCRβ functional identity was determined using V-family and J-gene, as those gave the most robust resolutions of TCRβ clones.

## Data Availability

All immunosequencing data underlying this paper can be downloaded and analyzed from Adaptive Biotechnologies’ immuneACCESS database at https://clients.adaptivebiotech.com/pub/Emerson-2017-NatGen

### Calculating Simpson Clonality

Simpson Clonality was defined as: 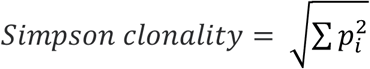 and was calculated on productive nucleotide rearrangements, where *p*_*i*_ is the proportional abundance of rearrangement *i* and *N* is the total number of rearrangements. Clonality values range from 0 to 1 and describe the shape of the frequency distribution. Clonality values approaching 0 indicate a very even distribution of frequencies, whereas values approaching 1 indicate an increasingly asymmetric distribution in which a few clones are present at high frequencies.

### Calculating a Generation Probability Cutoff for Common and Rare Clone Distinction

To calculate the generation probability, the TCRβ CDR3_aa_ sequence, and if available V family and/or J gene, of each clone were input into OLGA, an algorithm that calculates generation probabilities using a generative model for VDJ recombination (9). The unique rearrangements (a unique combination of TCRβ CDR3_aa_, V family, and J gene) from each sample were aggregated into a single data table and randomly shuffled. To optimize downstream computation efficiency, 31 sub tables (“chunks”) containing ∼2 million rearrangements were created. Within each sub table, rearrangements occurring greater than once were labeled as “common” and those occurring once labeled as “rare.” The generation probabilities of each sub table’s common and rare clones were plotted as density curves and their points of intersection identified. The median value of the 31 intersection points was used downstream as the universal cutoff point to identify individual rearrangements as either common or rare.

### Statistical Analysis for Association Between HLA Allele Similarity and TCRβ Clone Sharing

Mantel tests were conducted to evaluate the correlation between the pairwise number of shared HLA alleles and number of shared TCRβ clones. This correlation statistic is calculated by first creating two dissimilarity matrices, one for each variable being compared. The correlation between these two matrices is subsequently measured and then one matrix is repetitively shuffled to determine how often randomization of one matrix leads to increased correlation between the two matrices and thus variables. This permutation test evaluates the significance of the correlation between the observed dissimilarity matrices.

Linear mixed effects models were additionally created to measure the dependency of TCRβ clone sharing on HLA allele sharing by generating an intercept and slope. The mixed effects model format was selected to incorporate the impact of the numerous different sample comparisons between separate individuals resulting in random effects altering the regression.

All statistical analyses and visualizations were conducted using R version 3.6 (https://www.r-project.org/).

### Creating Null Distribution of TCRβ Clone Sharing Across groups

To test for the significance of the association between CMV or age with TCRβ clone sharing, we employed a permutation test in which the labels of interest were shuffled 1,000 times and the shuffled medians for each group were compared to the corresponding observed median. We then report the rank of each empirical median relative to the 1,000 shuffled medians and inclusive of the observed median, as well as the p-value for each group (S5A, S5B, S5C Figs). Rank values closer to 1000 signify that the observed median is greater than the permuted median in most or all of the permuations, while values closer to 0 indicate that the measured median was below most of the permuted medians. The p-value was determined as the number of permutations more extreme than the observed value plus 1, to be inclusive of the observed value, out of 1001 and multiplied by 2 to account for a two-sided test.

## ACKNOWLEDGEMENTS

We would like to thank all members of the Adaptive Biotechnologies Computational Biology Group who provided valuable insights during the project development and data analysis, in particular Erik Yusko, Bryan Howie, and Marissa Vignali.

## AUTHOR CONTRIBUTIONS

SAJ and SLS performed computational analysis and wrote the manuscript. RMG performed computational analysis and edited the text. JAR performed computational analysis. HSR and PAF supervised the study and edited the manuscript.

---

**Figure Supplement 1:**
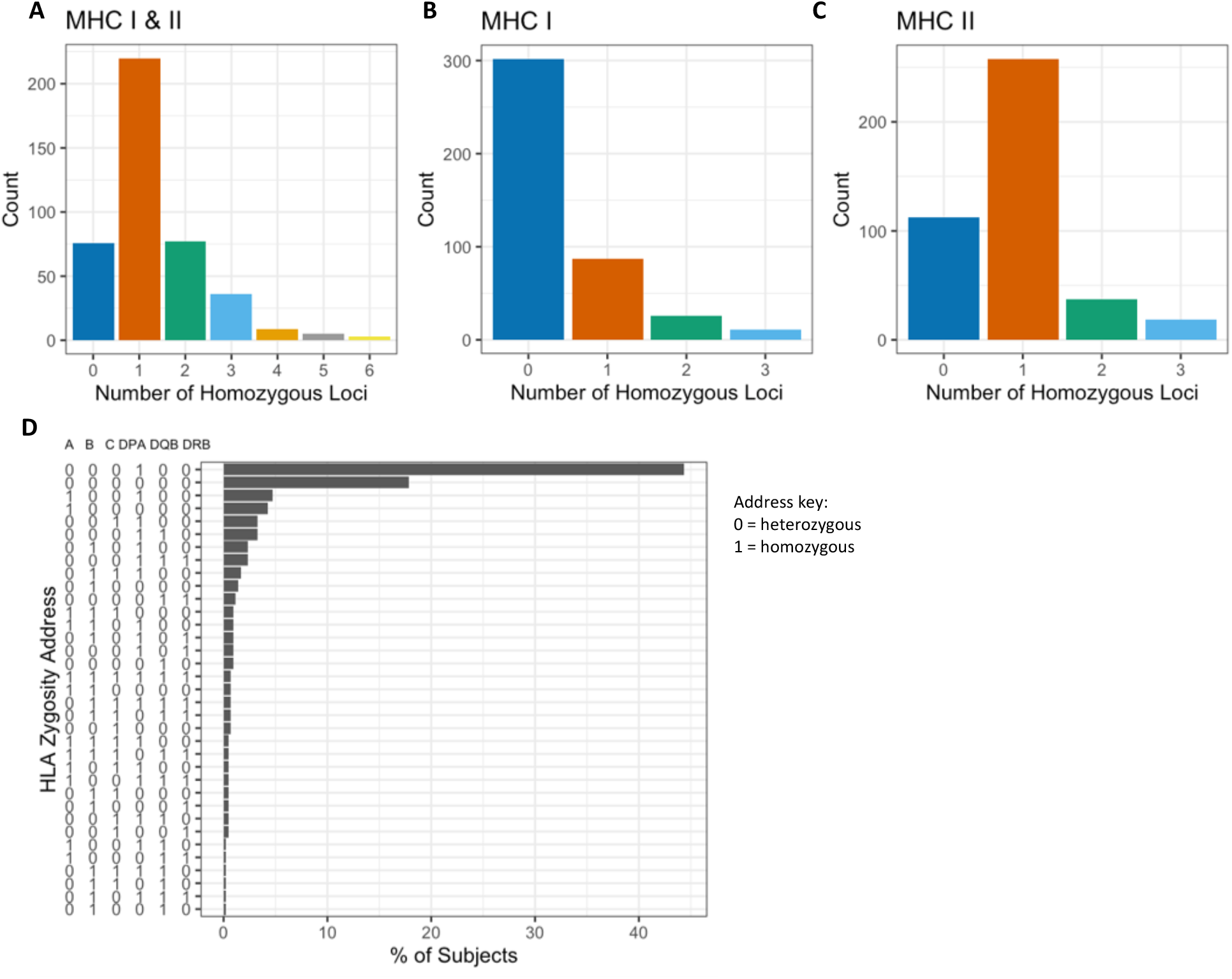
Distribution of HLA alleles in study cohort. (A) Count of individuals, among the cohort of 426, based on number of homozygous HLA loci. (B) Count of individuals based on number of homozygous HLA class I loci: HLA-A, HLA-B, HLA-C. Most individuals are heterozygous at all HLA Class I loci. (C) Count of individuals based on number of homozygous HLA class II loci: HLA-DPA1, HLA-DQB1, HLA-DRB1. (D) The distribution of individuals with each combination of homozygous loci. 44% of individuals in this cohort are homozygous at HLA-DPA1.

**Figure Supplement 2:**
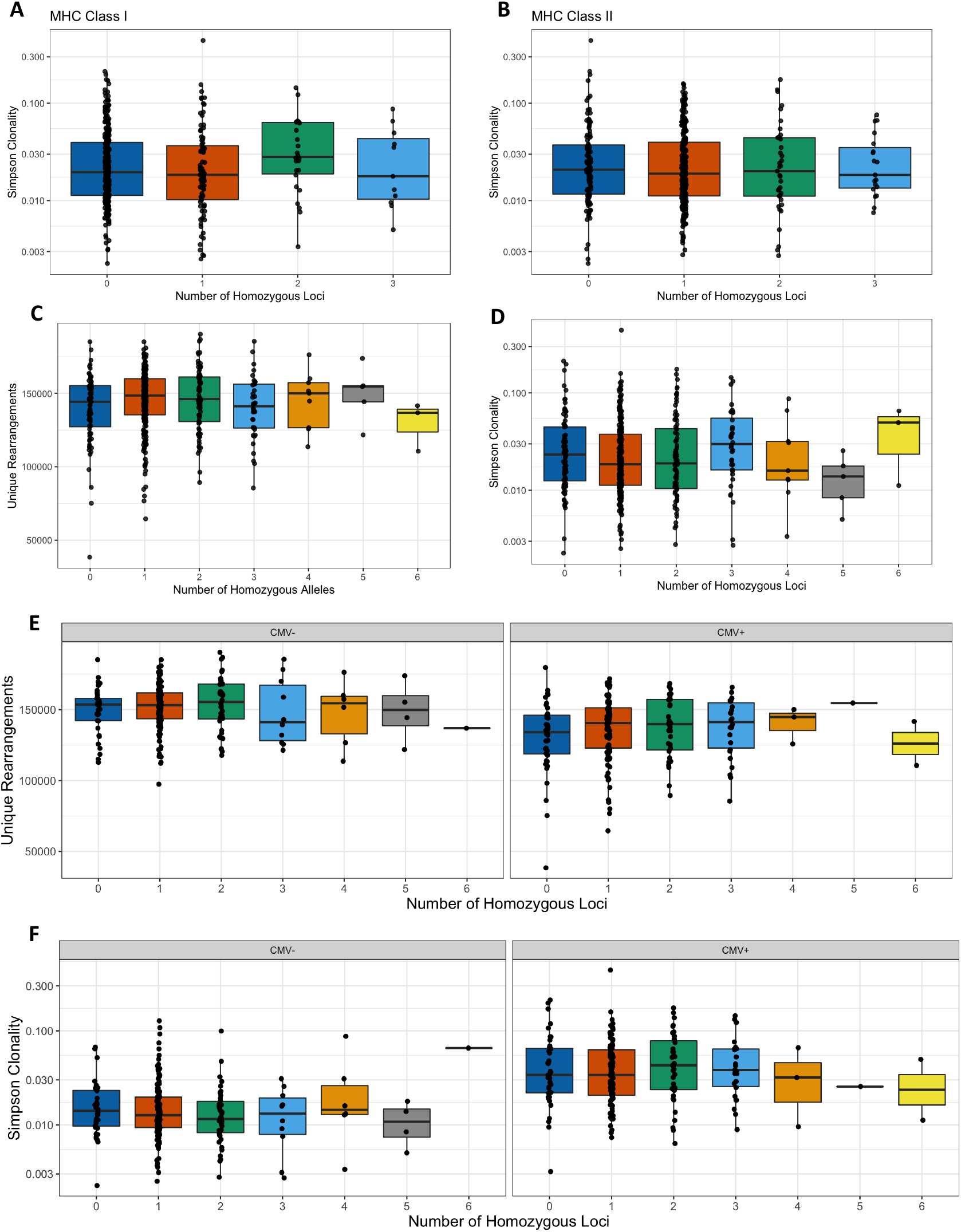
Impact of HLA zygosity on repertoire richness and clonality. HLA zygosity at neither (A) class I loci (Spearman rho = 0.011, p = 0.08) nor (B) class II loci were correlated with Simpson clonality (-0.023, p = 0.64). Overall, HLA zygosity in this cohort was not correlated with (C) richness (Spearman rho = 0.026, p = 0.60) or (D) Simpson clonality (Spearman rho = 0.0005, p = 0.99). (E) HLA zygosity was not correlated with richness among CMV-individuals (Spearman rho = 0.018, p = 0.79) or CMV+ individuals (Spearman rho = 0.086, p = 0.23). (F) HLA zygosity was not correlated with Simpson clonality among CMV-individuals (Spearman rho = -0.064, p = 0.34) or CMV+ individuals (Spearman rho = 0.023, p = 0.75).

**Figure Supplement 3:**
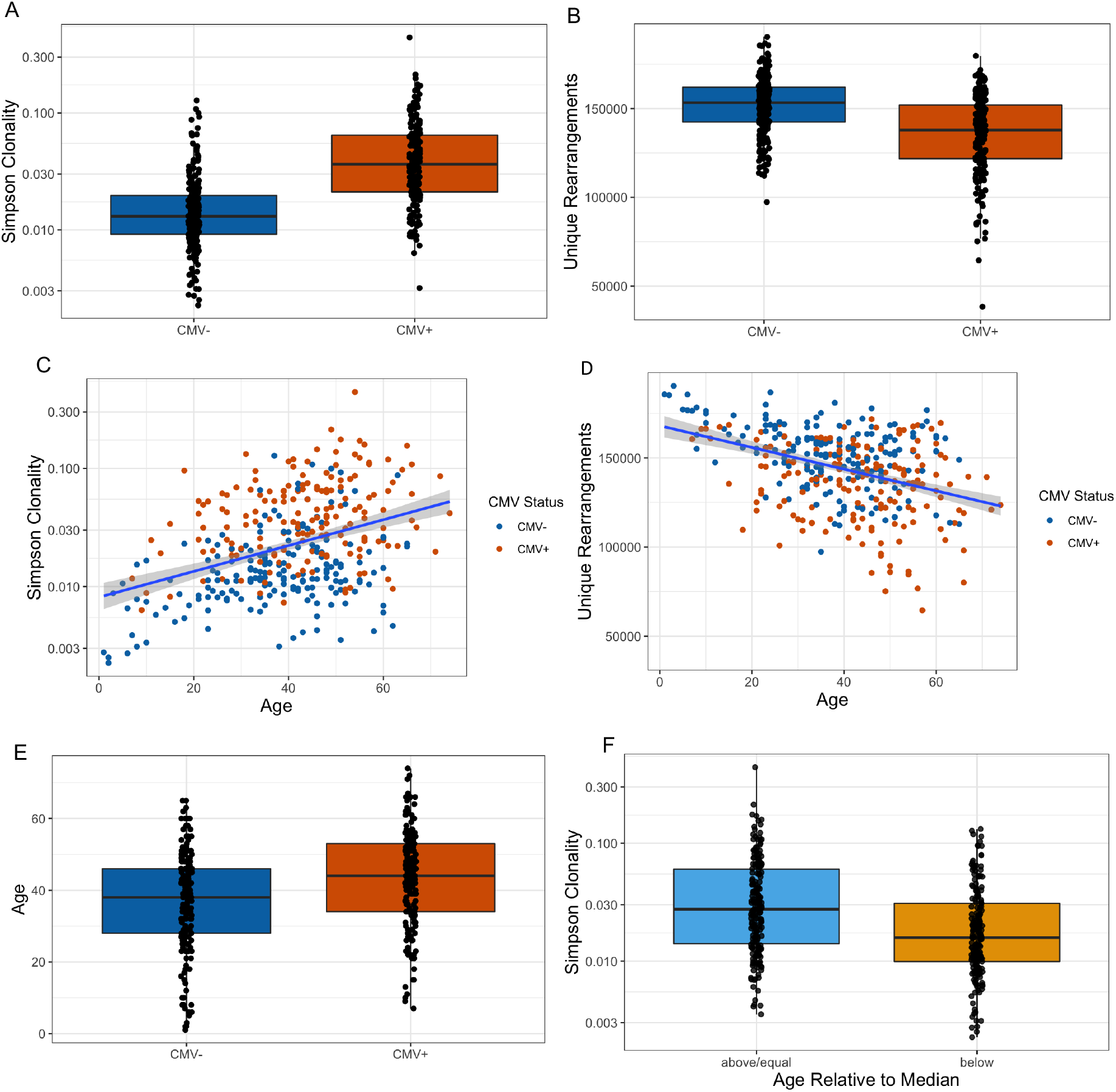
Impact of age and CMV status on repertoire clonality and richness. CMV+ Individuals had significantly (A) higher Simpson clonality (Wilcoxon rank sum test, p < 2.2e-16) and (B) lower richness (Wilcoxon rank sum test, p = 4.2e-14) than CMV-individuals. (C) Simpson clonality was positively correlated with age (Spearman rho = 0.34, p = 3.4e-11). (D) Repertoire richness was inversely correlated with age (Spearman rho = -0.35, p = 8.6e-12). (E) CMV+ individuals were significantly older than CMV-individuals (Wilcoxon rank sum test, p = 2.9e-5). (F) Subjects older than the overall median age had significantly greater Simpson clonality values than individuals that are younger than or equal to the median age (Wilcoxon rank sum test, p = 2.0e-7).

**Figure Supplement 4:**
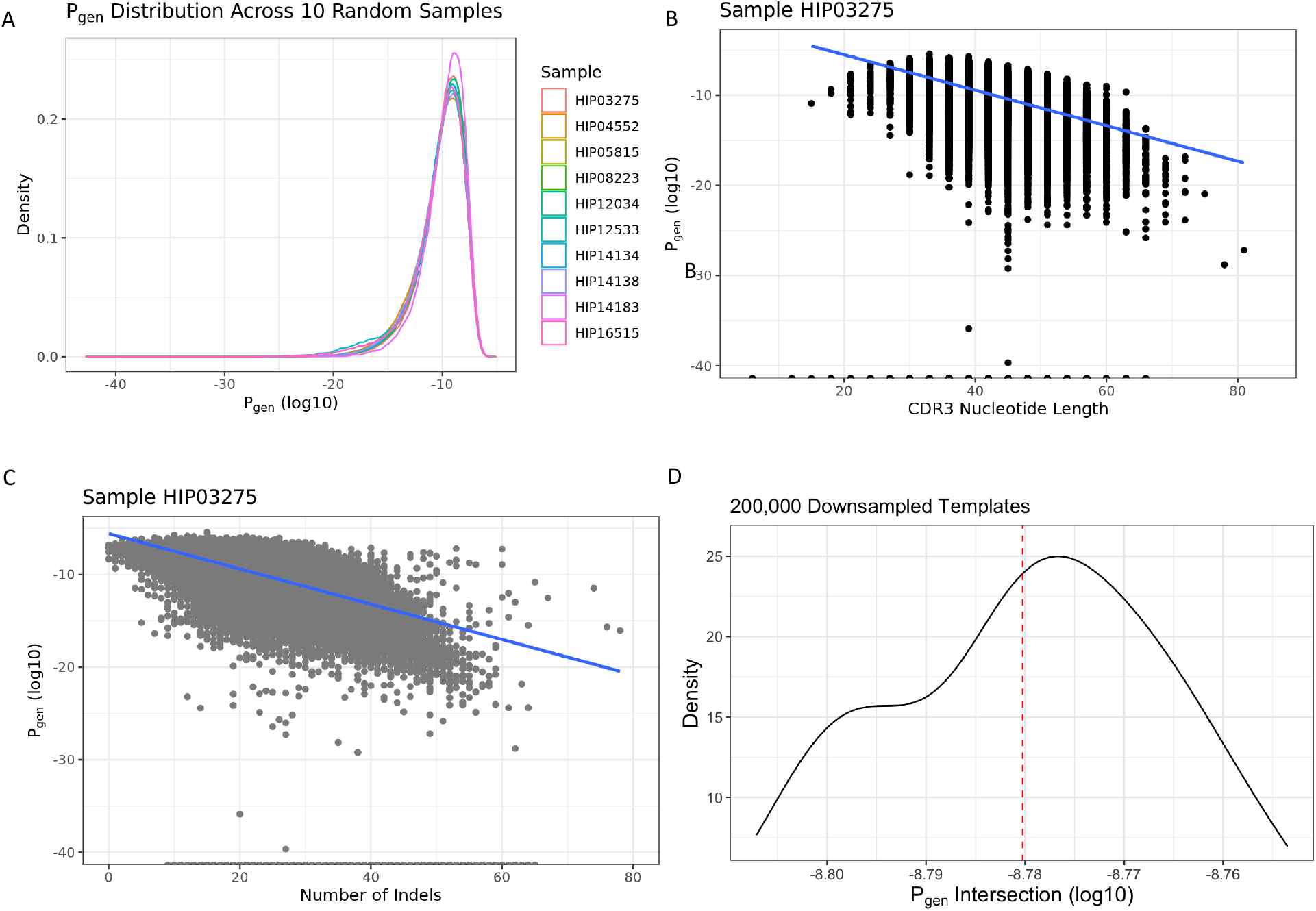
Distributions of TCRβ generation probabilities. (A) Distribution of generative probabilities of TCRβ clones from 10 representative individuals. Generation probability was significantly correlated with (B) CDR3 nucleotide sequence length (Spearman rho = -0.48, p < 2.2e-16) and (C) number of insertions and deletions (indels) (Spearman rho = -0.63, p < 2.2e-16), figure from single representative repertoire. (D) Identification of a generation probability cutoff point to distinguish between common and rare clones. Generation probability intersection points from the 31 chunks ranged from -8.81 to -8.76 (log10 transformed values) and the median -8.78 (1.66e-9) was selected as the cutoff point.

**Figure Supplement 5:**
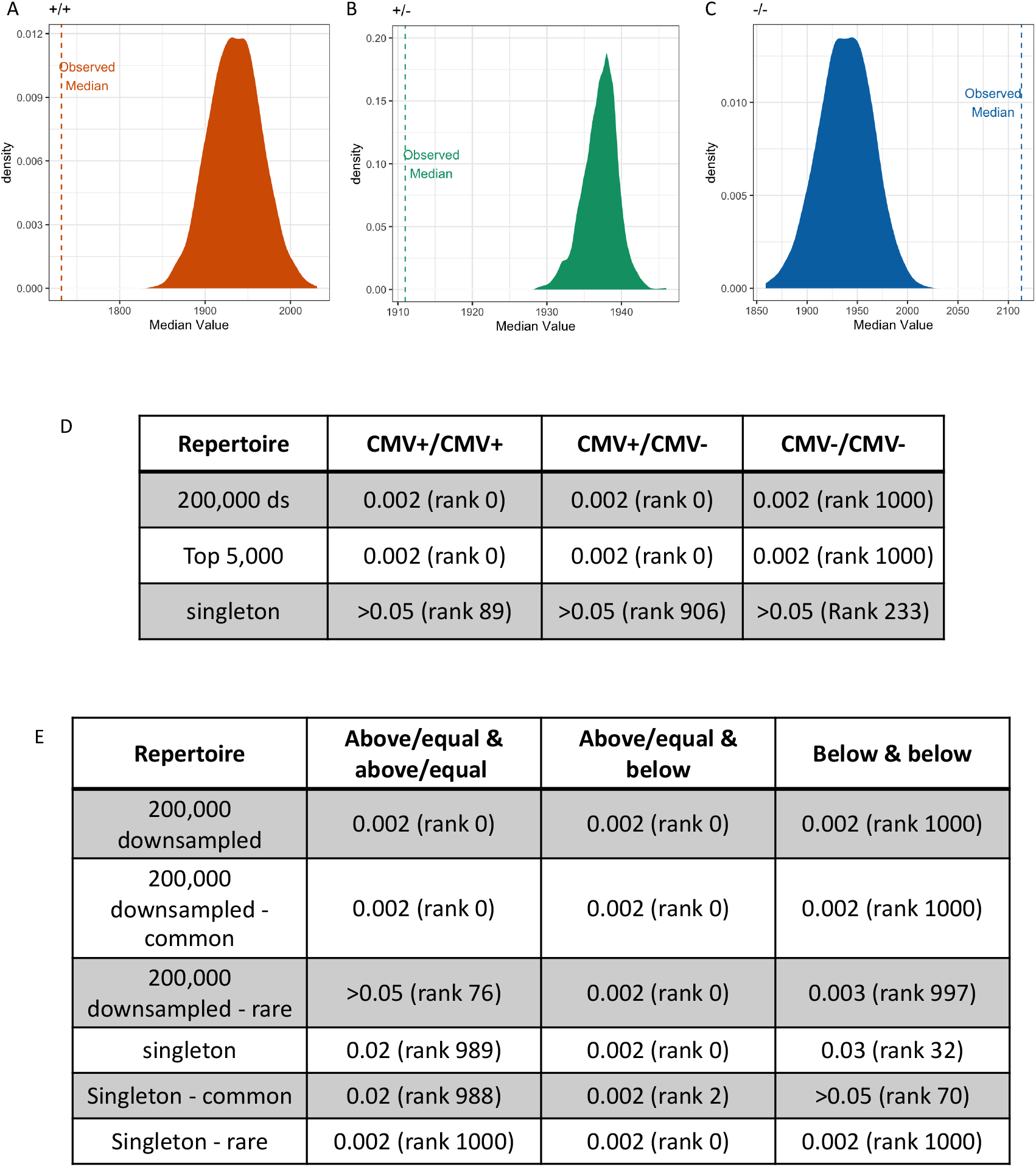
Determination of p-values by permutation tests. Within each set of pairwise comparisons, we compared the observed median value to the medians of 1,000 permuted comparisons and reported both rank and p-value for distribution indicated. (A) The observed median number of clones shared between CMV+ individuals was lower than the median of all shuffled comparisons, yielding a significant empirical p-value of 0.002. (B) The observed median number of clones shared between CMV+ and CMV-individuals was lower than the median of all shuffled comparisons, yielding a significant empirical p-value of 0.002. (C) The observed median number of clones shared between CMV-individuals was greater than the median of all shuffled comparisons, yielding a significant empirical p-value of 0.002. (D) Table containing the rank and empirical p-values of all clone sharing comparisons stratified by CMV serostatus. (E) Table containing the empirical p-values of all clone sharing comparisons stratified by age relative to median age (42).

**Figure Supplement 6:**
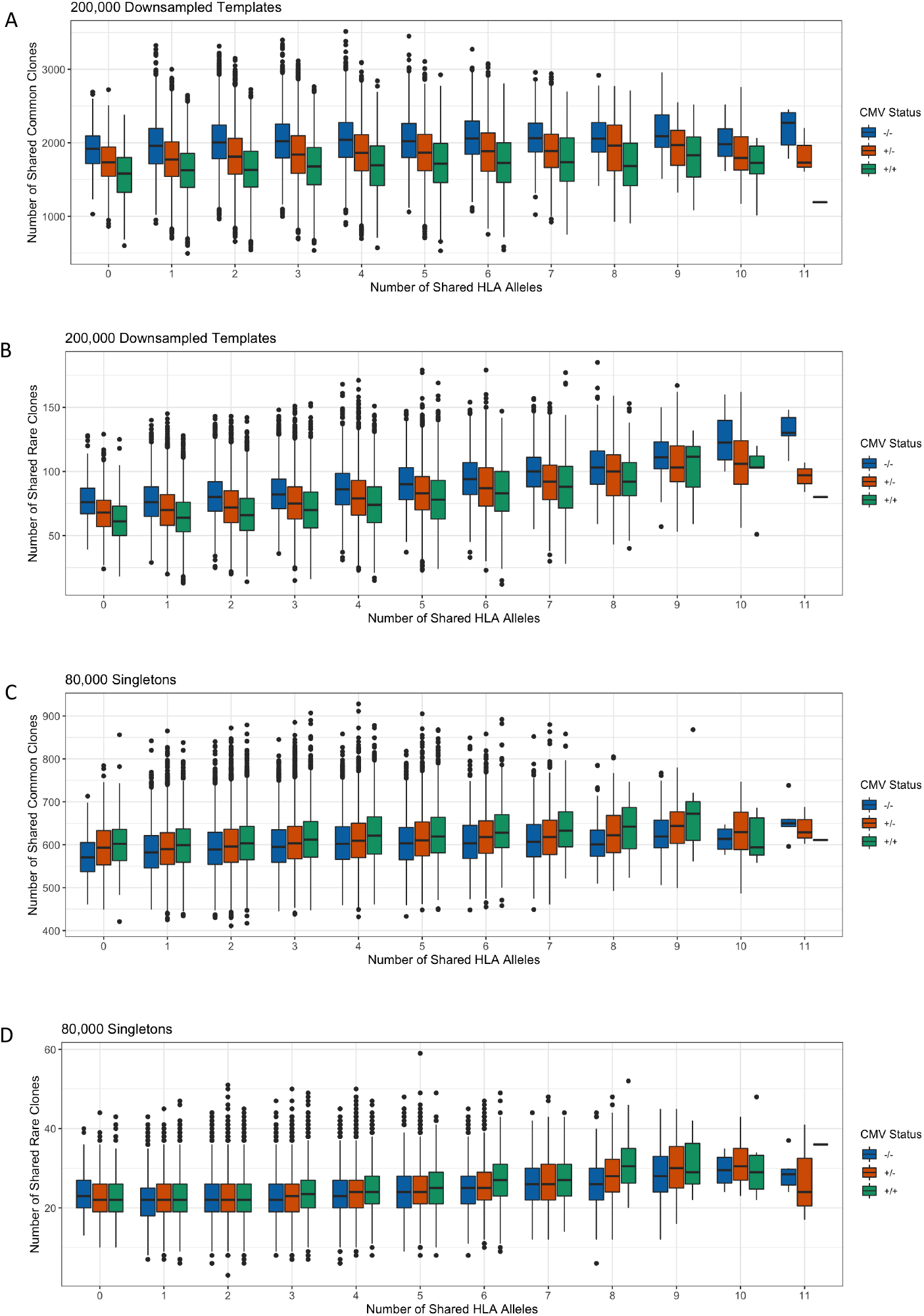
Impact of shared HLA alleles and CMV status on the sharing of common and rare TCRβ clones. (A) Individuals that were both CMV-shared more common clones than individuals that were both CMV+, regardless of the number of HLA alleles shared. (B) Individuals that were both CMV-shared more rare clones than individuals that were both CMV+, regardless of the number of HLA alleles shared. Individuals that were both CMV+ shared more (C) common and (D) rare singletons than individuals that were both CMV-, regardless of the number of HLA alleles shared.

**Figure Supplement 7:**
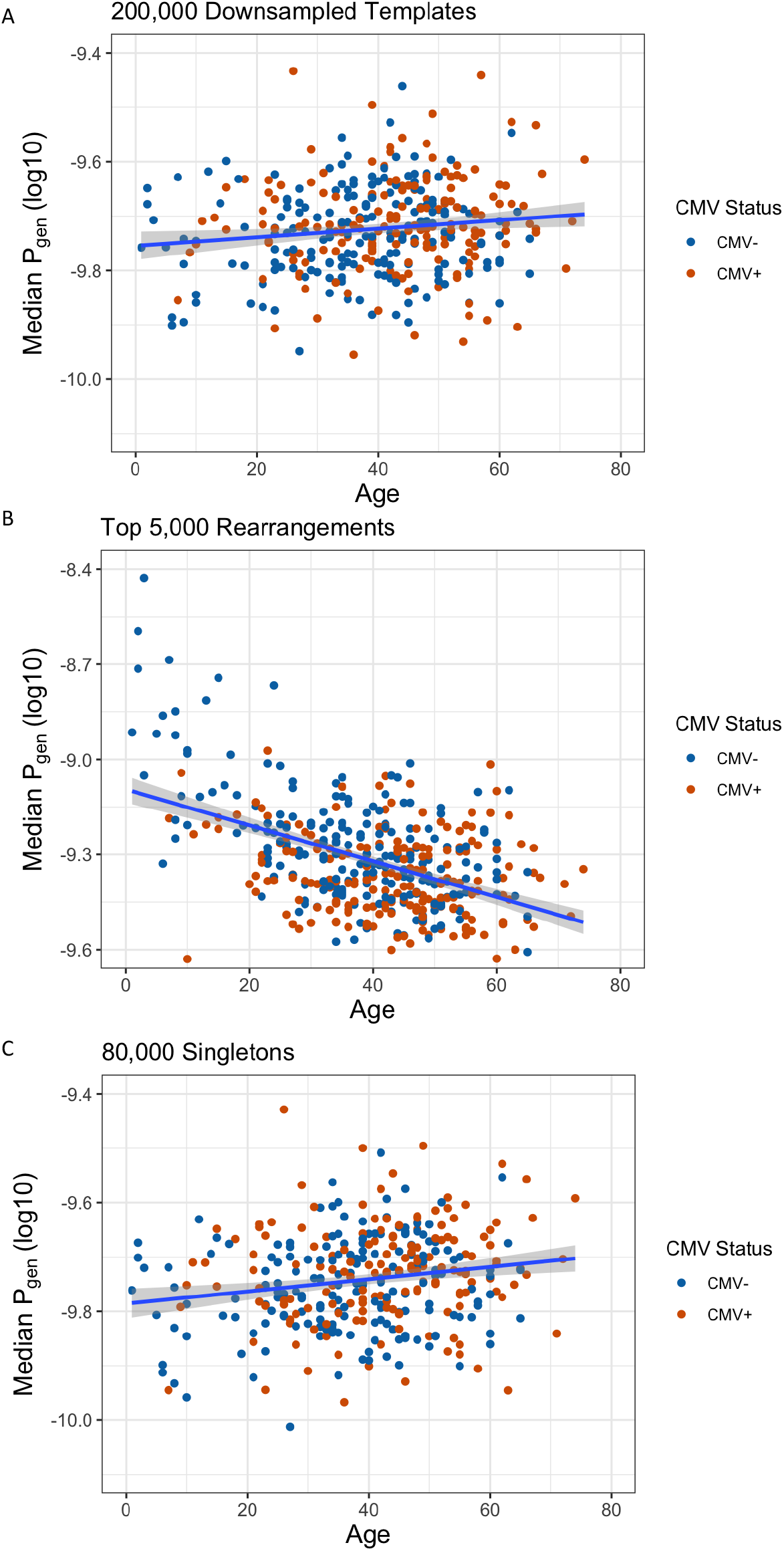
Association between age and median TCRβ repertoire generation probability. Age was significantly correlated with (A) higher median generation probability within the downsampled repertoires (Spearman rho = 0.12, p = 0.022), (B) lower median generation probability within the top 5,000 clones (Spearman rho = -0.37, p = 8.6e-15), and (C) higher median generation probability within the singleton repertoires (Spearman rho = 0.17, p = 0.0015). Only subjects with available age data were included in this analysis.

**Figure Supplement 8:**
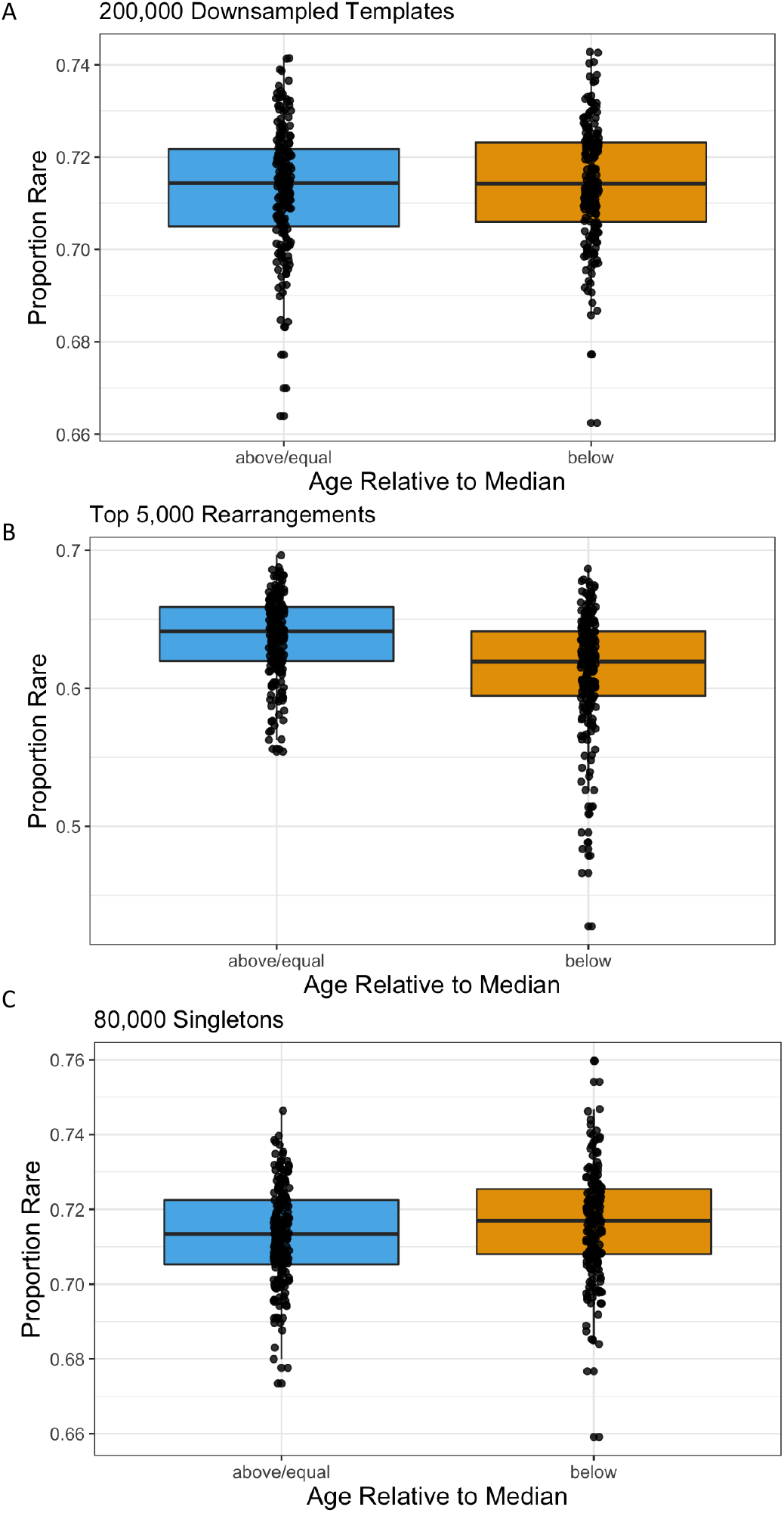
Proportion of rare TCRβ clones in repertoires. (A) There was not a significant difference in proportion of rare clones between older and younger individuals within the downsampled repertoires (Wilcoxon rank sum test, p = 0.5). (B) Older individuals had a significantly greater proportion of rare clones among their top 5,000 most abundant rearrangements than younger individuals (Wilcoxon rank sum test, p = 3.6e-10. (C) Older individuals had a significantly lower proportion of rare singletons compared to younger individuals (Wilcoxon rank sum test, p = 3.0e-2).

